# Space, Time and Episodic Memory: the Hippocampus is all over the Cognitive Map

**DOI:** 10.1101/150177

**Authors:** Arne D. Ekstrom, Charan Ranganath

## Abstract

In recent years, the field has reached an impasse between models suggesting that the hippocampus is fundamentally involved in spatial processing and models suggesting that the hippocampus automatically encodes all dimensions of experience in the service of memory. Here, we consider key conceptual issues that have impeded progress in our understanding of hippocampal function, and we review findings that establish the scope and limits of hippocampal involvement in navigation and memory. We argue that space and time serve as a primary scaffold to break up experiences into specific contexts, and to organize multimodal input that is to be associated within a context. However, the hippocampus is clearly capable of incorporating additional dimensions into the scaffold if they are determined to be relevant in the event-defined context. Conceiving of the hippocampal representation as constrained by immediate task demands—yet preferring axes that involve space and time—helps to reconcile an otherwise disparate set of findings on the core function of the hippocampus.

## “I am a traveler of both time and space, to be where I have been.” Robert Plant

Our understanding of the hippocampus, ranging from analysis of cellular properties up to its contributions to behavior, can be seen as one of the greatest success stories in modern neuroscience. Neuroscience research has tied the role of the hippocampus to spatial navigation, and there are compelling links between studies illustrating the properties of place cells, functional imaging, and neurophysiological recordings in humans during virtual navigation. Such findings provide considerable support for the idea of the hippocampus as representing a cognitive map of the current environment (O’keefe and Nadel, 1978).

There is also considerable evidence suggesting that the hippocampus contributes to a broad range of behaviors beyond spatial cognition. In rodents and primates, these include cells selective for odors, spatial views of landmarks, auditory cues, and conjunctions of task-specific variables, such as spatial context, landmarks, goals, and odors (Aronov et al., 2017; Ekstrom et al., 2003; O’Keefe and Speakman, 1987; Rolls and O’Mara, 1995; Wirth et al., 2017; Wood et al., 1999). In humans, there is near-consensus agreement that the hippocampus is necessary for episodic memory, and in particular, its contextual components (i.e., “when and where did I eat dinner last night?”) (Davachi and Dobbins, 2008; Eichenbaum et al., 2007; Ranganath, 2010). It is unclear, though, whether or how episodic memory is related to a broader “spatial” function within the framework of a cognitive map.

It is also clear that many extrahippocampal regions in humans and (to a lesser extent) in rodents, are sufficient to support spatial navigation and memory. For example, hippocampal lesions result in subtle deficits in spatial navigation in humans compared to the observed effects on episodic memory (Hirshhorn et al., 2012; Kolarik et al., 2016; Maguire et al., 2006). Lesions to areas outside of the hippocampus -- like precuneus, retrosplenial cortex, parahippocampal cortex, and posterior parietal cortex, appear to have much more profound affects on our ability to navigate (Barrash, 1998; Barrash et al., 2000; Mimori et al., 1998; Takahashi et al., 1997). Rodents with hippocampal lesions, while severely impaired at solving the Morris Water Maze (a classic task for navigational integrity), show some sparing of navigation, including inference of new navigational trajectories (Day et al., 1999; Pearce et al., 1998). Thus, space appears to involve a broader array of brain areas than the hippocampus, and this begs the question of what unique role the hippocampus plays in processing space more generally (Eichenbaum, 2017).

It is fair to say that we are at an impasse regarding the function of the hippocampus. Proponents of the cognitive map theory have used evidence from a wide range of tasks, ranging from scene perception to virtual navigation, and even prospective planning, to say that the hippocampus is preferentially or exclusively involved in “spatial processing” (Banta-Lavenex et al., 2014). The strongest version of this view that space is fundamental to all forms of cognition, and thus seemingly non-spatial tasks are really supported by spatial processing (Maguire and Mullally, 2013b). On the other side of the spectrum, evidence of “non-spatial” processing in the hippocampus, and extra-hippocampal contributions to space, have been used to refute the cognitive map theory of hippocampal function (Eichenbaum and Cohen, 2014). The strongest version of this view is that the hippocampus is involved in all aspects of declarative memory, and that space is just one of a nearly infinite set of variables that is automatically encoded by the hippocampus (Squire and Cave, 1991).

To move forward, we think it is useful to think about some basic questions that have gotten lost in the debate between the spatial and nonspatial perspectives and attempt to reconceptualize the function of the hippocampus more broadly in cognition. In this review, we will consider some significant questions about the role of the hippocampus in spatial and nonspatial cognition and we will review findings that establish the scope and limits of hippocampal involvement in navigation and memory. We argue that the hippocampus “maps” multimodal input in reference to a 4-D continuous topological representation involving space (3-D) and time (1-D). Critically, this mapping process is not rigid-- the specific hippocampal context map that is selected, the relative weighting of spatial and temporal dimensions, and the relevance of other “nonspatial” dimensions depend critically on behavioral relevance and task demands.

## What is the difference between “spatial” and “nonspatial”?

One key stumbling block in the ongoing debate about hippocampal function is that authors rarely clarify what they mean when they say that a behavior, task, or representation, is “spatial” or “nonspatial.” Although physicists can agree on a definition of space, most mammalian species do not have sensory receptors that can directly transduce information about space. As pointed out centuries ago by Immanuel Kant, space must be constructed in relation to a reference frame. Depending on how we navigate, we can consider the positions of objects in relation to our immediate peripersonal space, our larger, unseen environment (i.e., the town we are in), or even larger geographic context (i.e., the state or country we are in), which we may never actually navigate but can readily see on a map (Montello, 1993). Objects typically have a position in space, which we can define relative to our current body position (termed “egocentric”), or relative to one another (termed “allocentric”). Space also can also be topological, rather than Cartesian, meaning that it can preserve the relative, rather than absolute positions of objects to each other (Poucet, 1993).

Tolman proposed that we construct a cognitive map of space, which refers to a metric, viewer-independent representation of the locations of objects (Tolman, 1948). O’Keefe and Nadel (1978) subsequently argued that the hippocampus accomplishes this basic function via neurons that fire at specific spatial locations during navigation (i.e., “place cells”). O’Keefe and Nadel’s (1978) monograph provides a detailed account of the cognitive map model, but most empirical studies of hippocampal function do not articulate such precise hypotheses. This state of affairs has resulted in a tendency for circularity, such that spatial processing is operationally defined in reference to the variables that happen to be related to hippocampal involvement in a task (Bird and Burgess, 2008; Hassabis and Maguire, 2007). For example, some studies have found that the hippocampus shows increased activity during perception, encoding, imagery, or working memory maintenance of scenes (Aly et al., 2013; Barense et al., 2010; Bird et al., 2010; Hassabis et al., 2007a; Preston et al., 2010; Stern et al., 2001; Summerfield et al., 2010), and patients with hippocampal damage appear to be deficient at imagining scenes (Hassabis et al., 2007b) (but see Hurley et al., 2011; Mullally et al., 2014, for conflicting findings). Based on such findings, Maguire and colleagues (Maguire and Mullally, 2013a; Mullally and Maguire, 2014) proposed “scene construction theory” (“SCT”), which argues that, “….the hippocampus primarily acts to facilitate the construction of atemporal scenes and in doing so allows the event details of episodic memories and imagined future experiences a foundation on which to reside. In this way, hippocampal-dependent scene construction processes are held to underpin and support episodic memory, predicting the future, spatial navigation, and perhaps even dreaming and mind-wandering…” (Mullally and Maguire, 2014).

Studies of scene encoding and imagery are often seen as supporting the Cognitive Map theory. Moreover, SCT is often taken as a modern version of the Cognitive Map Theory, in that it attempts to accommodate human episodic memory and prospective thinking within the broader umbrella of spatial processing that is fundamentally supported by the hippocampus. However, in its attempt to accommodate a wide range of behaviors, scene construction theory flexibly reframes space in a manner that is not obviously consistent with the original idea of the cognitive map. Whereas Cognitive Map Theory specifically refers to the idea that the hippocampus supports a metric, allocentric representation of space, there is no reason to assume that scene perception, recognition, or imagery should require an allocentric representation.

Scenes are routinely considered as “items” in memory studies (Canli et al., 2000; Ryan et al., 2000), and in many studies, scenes include multiple objects. Scenes are also typically experienced egocentrically, and there is no requirement that they maintain metric properties during encoding, retrieval, or visual imagery. For these reasons, Burgess, Maguire, & O’Keefe (2002) stated, “A fundamental distinction exists between simple iconic representations of single objects or 2D scenes and representations that include knowledge of the 3D locations of the elements of a scene. For example, either might suffice for recognition of a scene as familiar, but the latter would be needed to decide upon a novel shortcut or appreciate what the scene would look like from another point of view.” Burgess et al. (2002) went on to expand on Cognitive Map Theory by proposing that, “the parahippocampus supports processing of the spatial information present in visual scenes… [and] the hippocampus appears only to be activated in more complex navigational situations”.

Of course, SCT does not have to be consistent with Cognitive Map Theory, but as a theory, SCT should be falsifiable. Unfortunately, to our knowledge, no version of SCT has ever defined the characteristics of a scene. Scenes are by no means a homogenous category, and a picture of a natural scene can differ in countless ways from a picture of an object. Given that SCT does not specify what the hippocampus does during scene construction, and that it does not provide an explicit definition of a scene, it is not clear that SCT is a viable or falsifiable theory of hippocampal function. More generally, we see the lack of concern about representation as a significant issue in the interpretation of research linking the hippocampus to perception, encoding, and imagination of scenes. There is certainly evidence linking the hippocampus to processing of scenes (e.g., Aly et al., 2013; Barense et al., 2010; Bird et al., 2010; Hassabis et al., 2007a; Preston et al., 2010; Stern et al., 2001; Summerfield et al., 2010), but we cannot understand the significance of this evidence in the absence of well-articulated hypotheses about how the hippocampus represents information in scenes. One idea that potentially bridges scene processing with the Cognitive Map Theory is that the hippocampus could encode information in scenes in reference to a location in a spatial context (Georges-Francois et al., 1999), and we will explore this idea further in the next section.

## How is space mapped by the hippocampus?

When considering exactly what the hippocampus does, it is useful to think about the characteristics of place cells. Although there are good reasons to draw analogies between place cells and a cognitive map, overwhelming evidence suggests that place cells do <u>not</u> map space in a manner that is akin to a GPS. For instance, many studies have shown that when a rat runs on a linear track, place cells remap (i.e., the place fields for individual cells differ) when a rat runs in one direction compared to another (Gothard et al., 1996; Redish et al., 2000). Perhaps more fundamentally problematic for the idea of a global Cartesian map of space, place cells also change their preferred location of firing depending on the future goal of the animal (Ainge et al., 2007; Ferbinteanu and Shapiro, 2003; Lee et al., 2006; Wood et al., 2000). When an animal navigates toward a goal location on a T-maze, for instance, different place cells fire depending on whether the goal is on the right or left side of the maze. The results from T-maze and linear track paradigms demonstrate that the same location in space can be referenced to competing hippocampal place cell maps, depending on the behavioral context. Similar findings have also been shown in humans: place cells change where they fire depending on the store that a patient is searching for in virtual reality, and also fire to conjunctions of viewing landmarks, searching for goals, and spatial locations (Ekstrom et al., 2003). These findings make clear that the hippocampus does not indicate precise cartographic location within the world. Instead, it indicates topologically accurate coordinates within a multidimensional context space, which are shaped strongly by the ability of the animal to infer the variables necessary for solving the task.

Even studies of place cell activity during simple random foraging reveal that hippocampal place cell maps are anchored to specific spatial contexts (Muller and Kubie, 1987; Skaggs and McNaughton, 1998; Wilson and McNaughton, 1993). Place cells can indicate the location of an animal in a particular spatial context, but the same cells would have different place fields in a different, nearby context. This raises an interesting question-- if place cell codes are bound to specific spatiotemporal contexts rather than global spatial coordinates, how can the hippocampus map relative locations across context boundaries? For instance, if you ring your neighbor’s doorbell, and you hear her walk to the door, you might know that she is in close proximity. However, the hippocampal place cell code could not be used to figure this out, because she is outside of your current context. Otherwise, there would be immediate interference between the current location mandated by the hippocampus and the imagined locations. Moreover, it is not clear that the hippocampal place code could be used to determine that your house is closer to your neighbor’s house than to a house on the opposite side of the planet.

Our example illustrates the point that that the hippocampus does not necessarily represent a global cognitive map, but rather that it represents a series of maps, each tied to a different spatial context. In the case of the T-maze and linear track paradigms, competing maps can even be tied to the same spatial context (see also McKenzie et al., 2014; McKenzie et al., 2013). Critically, if we conceptualize the hippocampal place code as contextually-driven, rather than globally accurate, we solve the problem of otherwise catastrophic interference of competing place cell maps. This point accords with an idea that is well established in the human spatial navigation literature -- we often construct representations of disconnected “microenvironments” without updating our position relative to the global “macroenvironment” (Han and Becker, 2013; Wang and Brockmole, 2003a; Wang and Brockmole, 2003b). In other words, when we navigate in a building, we encode the locations and features that are useful for remembering locations within the building, with little memory or knowledge of how the landmarks and directions we have chosen relate to outdoor locations or other buildings we have recently explored. The idea that a microcontext -- be it a spatial goal or nearby environment that we explore, can remain unconnected with other neighboring contexts is entirely consistent with findings suggesting goal and context remapping in the hippocampus. Rather than providing a systematic map of known space, the hippocampus constructs multiple maps with overlapping content, and the selection of the appropriate map is based on many factors beyond space. In the next section, we focus on the most important factor—time (Ranganath and Hsieh, 2016).

## What does cognitive mapping have to do with episodic memory?

Just as there is a general consensus that the hippocampus is critical for allocentric spatial memory in the rodent brain, there is near consensus that it is critical for episodic memory in the human brain. Virtually every paper on hippocampal representation of space begins with a perfunctory reference to episodic memory (e.g., Bird and Burgess, 2008) (Burgess et al., 2002; Buzsaki and Moser, 2013). Buzsaki & Moser (2013) went so far as to equate path integration with episodic memory (despite the fact that the very idea of “integration” implies something greater than a single episode). In most instances, however, the exact relationship between a cognitive map and episodic memory is left to the reader’s imagination. This is probably because it is not clear how a purely spatial code would be sufficient to support episodic memory tasks, such as remembering where you left your glasses *last night*, or where you parked your car *this morning.* These memories would be nearly impossible to retrieve due to interference with countless other memories for the locations of your glasses and keys.

O’Keefe & Nadel (1978), considered this issue, and in accord with Tulving’s (1972) definition of episodic memory, they stated that the hippocampus supports, “memory for items or events within a spatio-*temporal* context.” Studies of hippocampal “time cells” are consistent with this idea (Eichenbaum, 2014). Much as a population of place cells provides a map of a spatial context, a population of time cells maps the progression of time through a situational context. More specifically, time cells fire at specific time points as an animal experiences a predictable sequence of events, even when the animal remains in the same location (Kraus et al., 2015; MacDonald et al., 2013; MacDonald et al., 2011; Naya and Suzuki, 2011; Pastalkova et al., 2008; Wang et al., 2015). Moreover, just as place cells remap when an animal is moved to a new spatial context, hippocampal time cells “retime” when the temporal structure of a task, or the current behavioral context is changed (MacDonald et al., 2013; MacDonald et al., 2011; Sakon et al., 2014). These findings demonstrate that hippocampal time and place cells “map” experiences in relation to a context that is bounded in time and space. Consistent with the data from rodents, our labs have repeatedly shown that activity patterns in the human hippocampus carry information about the spatiotemporal context in which an object was encountered (Dimsdale-Zucker et al., 2017; Ekstrom et al., 2011; Hsieh et al., 2014; Kyle et al., 2015; Libby et al., 2016; Lieberman et al., 2017; Zhang and Ekstrom, 2013).

With our current understanding of time cells, we can see how the hippocampus might represent experiences as they unfold within the boundaries of an episode. But what is an “episode”? Research on event segmentation provides one answer to this question. Zacks and colleagues have shown that, when watching a movie or reading a narrative, people tend to parse temporally continuous information into discrete events (Zacks and Swallow, 2007). Just as a large border (such as a wall) leads one to assume a boundary between two spatial contexts, large changes in the continuity of incoming information over time appear to trigger boundaries between events (Speer and Zacks, 2005; Zacks et al., 2001). Interestingly, both event boundaries (Axmacher et al., 2010; Baldassano et al., 2016; Ezzyat and Davachi, 2014; Hsieh et al., 2014) and spatial boundaries (Bird et al., 2010) modulate hippocampal activity. These studies demonstrate how hippocampus often uses temporal information, along with spatial information, to represent task-specific aspects of context (Watrous and Ekstrom, 2014).

Putting together what we have reviewed so far, we propose that the hippocampus defines contexts as chunks of experience that have low variance across space and time (Fuhs and Touretzky, 2007; Zucker and Ranganath, 2015). When the variance across experiences exceeds a threshold, we form a new context by differentiating unique temporal epochs (Fuhs and Touretzky, 2007). According to this view, we can expect the hippocampus to treat any cue that remains stable over a chunk of time spent in a given place as an indicator of the spatial context, and sufficient change in one of these cues can lead one to infer the existence of a new context. As an example, if the configuration of furniture remains the same every time you walk into your office, then those factors will become indicators of the context (or “landmarks”), and hippocampal cells would likely map your ongoing experiences relative to the boundaries of your office (Figure 1a). If, one day, the furniture configuration has changed in your office, the change would trigger the hippocampus to assign distinct context representations to the “old” vs. “new” office configurations (Figure 1b). Additionally, if some pieces of furniture, such as your office chair, change locations continuously, these would be unlikely to serve as indicators of your current spatial context. Instead, the chair would be treated as an item to be mapped in the office context (Figure 1c).

**Figure 1.**
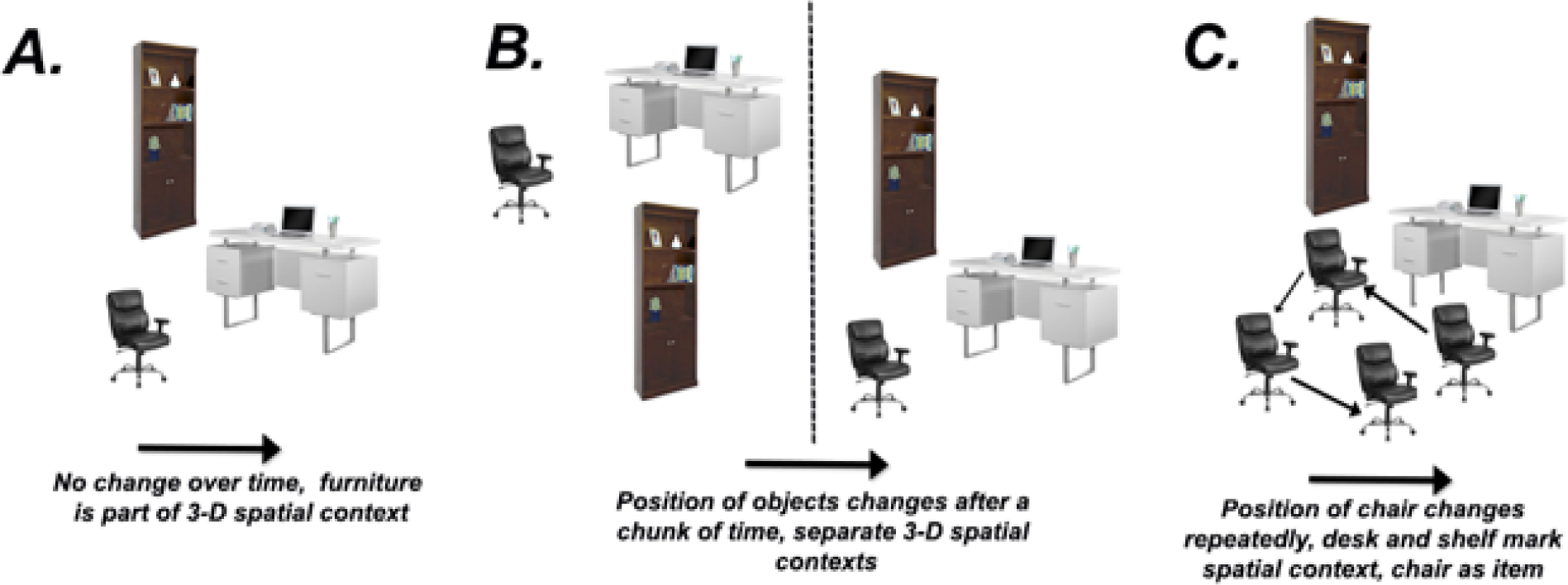
Factors that can influence hippocampal representations of spatial contexts.

## When does hippocampus go beyond space and time?

Until now, we have presented a view in which the hippocampus maps people and things to spatial and temporal positions within the boundaries of an event context (Eichenbaum, 2013; O’keefe and Nadel, 1978; Zucker and Ranganath, 2015). The implication is that hippocampal maps are limited to four dimensions (3-D space +1-D time). Recent studies, however, have shown that hippocampal cells encode dimensional information that extends beyond space and time--as just one example, Aronov et al. (2017) trained rats to perform complex mappings between sounds and manual responses (Aronov et al., 2017). They found that hippocampal cells formed discrete firing fields at particular sound frequencies. These findings, along with findings of selectivity for other nonspatial features (e.g., odors) clearly indicate that the hippocampus can add dimensions beyond space and time to the cognitive map. Functional imaging studies in humans have also indicated that the hippocampus can encode abstract spaces such as social hierarchies (Tavares et al., 2015). Does this mean that time and space are no different than any other variable? Should we assume that the hippocampus simply assigns dimensional axes to all continuous variables, such that a hippocampal event representation has almost infinite dimensionality?

An alternative way of thinking about the issue relates back to our earlier discussion. The hippocampus has a strong bias to encode features within a 4-D spatiotemporal framework, but it also encodes variables that are behaviorally relevant, given the current context. Space and time have a privileged status because they are used to define and learn about the current context. For example, rats in the Aronov et al. experiment were trained to perform a “sound manipulation task” during sustained chunks of time in an operant conditioning chamber. We think it is reasonable to speculate that hippocampal cells encoded information about the spatial context and the temporal structure of task trials from the very beginning, whereas auditory selectivity became apparent only after the animal learned that sound frequencies are behaviorally relevant in the context. The critical point is that the hippocampus uses regularities in the environment across time and space in order to define a context (Eichenbaum, 2013; O’keefe and Nadel, 1978; Ranganath and Hsieh, 2016), and this initial representation must be established in order to learn what is behaviorally relevant in that context. Once the behaviorally relevant variables are determined, then these dimensions are added to the cognitive map.

Although laboratory rats with limited experience of the world must rely on gradual learning to incorporate new dimensions in the hippocampal map, we suspect that, in adult humans, the process is likely to be much faster. To understand why this might be the case, it is helpful to consider the information that gets *in* to the hippocampus. Considerable evidence suggests that the hippocampus, along with the diencephalon, closely interacts with a posterior medial network that includes the parahippocampal, retrosplenial, and medial entorhinal cortex (Bergmann et al., 2016; Kahn et al., 2008; Libby et al., 2012; Maass et al., 2015). The posterior medial network, like the hippocampus, is extensively engaged in spatial navigation and episodic memory. Ranganath & Ritchey (Ranganath and Ritchey, 2012; Ritchey et al., 2015) proposed that the posterior medial network encodes schemas that specify the gist of the spatial, temporal, and causal relationships that apply within a particular event context. Similar proposals regarding a network basis for spatial navigation have also been proposed (Ekstrom et al., 2014; Ekstrom and Watrous, 2014). There is considerable evidence suggesting that ventromedial prefrontal cortex, possibly via the nucleus reuniens (Ito et al., 2015), plays a critical role in regulating hippocampal encoding of information beyond space and time (Gruber et al., 2017; Navawongse and Eichenbaum, 2013; Place et al., 2016; Young and Shapiro, 2011). Notably, the ventromedial prefrontal cortex is extensively interconnected with the posterior medial network, and prefrontal-hippocampal interactions may be critical for both incorporating new information into neocortical event schema representations (Preston and Eichenbaum, 2013; van Kesteren et al., 2012; Zeithamova and Preston, 2010).

The more general point is that the hippocampus is situated between large networks of brain regions, and its role in cognition can be dynamically configured dependent on the relevant extra-hippocampal brain areas with which it interacts. Interactions with neocortical network “hubs” might be a mechanism for the hippocampus to preferentially emphasize any dimension of information coding. In this way, extrahippocampal cortical brain hubs specify the dimensions beyond space and time that are relevant for representation by the hippocampus (Schedlbauer et al., 2014; Zhang and Ekstrom, 2013).

## Closing thoughts

At the outset of this paper, we stated that the field has reached an impasse between two views of the hippocampus--one that places spatial mapping as its primary function and another that sees the hippocampus as indiscriminately mapping all dimensions of experience. Both of these views capture essential points--any viable theory of hippocampal function needs to account for the critical role of the hippocampus in spatial cognition and its role in episodic memory (Schiller et al., 2015). Spatial and temporal information are used to break up our experiences into specific events that occurred at specific places (Eichenbaum, 2013; O’keefe and Nadel, 1978; Ranganath and Hsieh, 2016). Although the hippocampus uses space and time as a primary scaffold for defining contexts, and for organizing incoming information within a context representation (O’keefe and Nadel, 1978), other dimensions can be incorporated into the scaffold (Eichenbaum and Cohen, 2014) if they are determined to be relevant in the event-defined context. Conceiving of the hippocampal representation as constrained by immediate task demands -- yet preferring axes that involve space and time -- helps to reconcile an otherwise disparate set of findings on the core function of the hippocampus.

## Acknowledgements

ADE wishes to acknowledge funding from NIH/NINDS grants NS076856, NS093052 (ADE), and NSF BCS-1630296 and CR acknowledges funding from a Vannevar Bush Fellowship (Office of Naval Research Grant N00014-15-1-0033). Any opinions, findings, and conclusions or recommendations expressed in this material are those of the author(s) and do not necessarily reflect the views of the Office of Naval Research or the U.S. Department of Defense.

